# X-ray crystal structure of the N-terminal domain of *Staphylococcus aureus* cell-cycle protein GpsB

**DOI:** 10.1101/2025.08.05.668015

**Authors:** Nathan I. Nicely, Thomas. M. Bartlett, Richard W. Baker

## Abstract

GpsB is a conserved cell-cycle regulator in the Firmicute clade of Gram-positive bacteria that coordinates multiple aspects of envelope biogenesis. Recent studies demonstrate interactions between GpsB and the key division cytoskeleton FtsZ, suggesting that GpsB links cell division to various aspects of cell envelope biogenesis in Staphylococcus aureus and potentially other Firmicutes. We determined a 1.7 Å resolution crystal structure of the N-terminal domain of *Staphylococcus aureus* GpsB, revealing an asymmetric dimer with a bent conformation. This conformation is nearly identical to one of two conformations reported by Sacco, et al., confirming the unique conformation of *S. aureus* GpsB compared to other gram-positive bacteria. This structural agreement provides strong validation of the *S. aureus* GpsB fold and supports its proposed role in organizing the cell division machinery.

## Description

GpsB is a widely conserved adaptor protein in Gram-positive Firmicutes (synonym Bacillota) that coordinates cell-cycle progression by coupling cell envelope biogenesis to the cell-division machinery. GpsB has two structured domains — an N-terminal dimerization domain and a C-terminal trimerization domain — which are separated by a flexible linker^1,2^. Thus, GpsB is thought to act as a hexamer that can bind to multiple cell division proteins simultaneously, thereby facilitating the spatiotemporal coordination of various molecular machines. The N-terminal domain of GpsB, in addition to containing a membrane-binding loop that drives binding to the inner leaflet of the membrane^1^, also binds small peptides bearing a consensus (S/T)-R-X-X-R−(R/K) motif. This motif is found in proteins such as PBP1A^3^, FtsZ^4,5^, EzrA^6^, DivIVA^7^, TarO/G^5^, and FacZ^8^, allowing GpsB to coordinate division site localization of multiple protein complexes. Consistent with these binding motifs, GpsB directly binds and regulates FtsZ^4^, contributing to the correct placement of FtsZ and other Staphylococcal division proteins^8^ and coordinating cell envelope growth to division^5^, underlining its intriguing potential to communicate information between cell envelope synthesis and morphogenetic factors. Unsurprisingly, and presumably as a result of its coordination of these features, GpsB is necessary for normal *S. aureus* morphogenesis^9,10^. GpsB’s interactions with FtsZ^11^ and various cell envelope synthesis factors^3^ are conserved in other Firmicutes, suggesting its role as an adaptor between cell cycle and envelope growth is also broadly conserved.

Prior to 2024, structures for the N-terminal domain of GpsB had been described for *Bacillus subtilis*^3,12^, *Listeria monocytogenes*^3^, and *Streptococcus pneumoniae*^3^. These GpsB orthologs adopt a conserved fold, showing a long parallel two-helix bundle with two short helices that form a 4-helix ‘cap’ at one end^1^. Notably, the central helical bundle is a rigid helix, reinforcing the model of GpsB as a linear adaptor scaffold.

Recently, Sacco et al. revealed a novel conformation of *S. aureus* GpsB (*Sa*GpsB)^13^, where the N-terminal homodimer adopts an asymmetric dimer, in which two protomers display a kinked helix conformation, mediated by a hinge formed by a three-residue insertion exclusive to Staphylococcus species. This hinge comprises a cluster of methionine residues (“MAD” or “MNN” insertion) not found in other Firmicutes, conferring conformational flexibility. Excising this insertion increases thermal stability and abolishes an overexpression lethal phenotype in *Bacillus*, suggesting functional tuning via flexibility. Thus, functional and structural divergence appears between *S. aureus* and other Gram-positives. Whereas GpsB in other species is rigid, *Sa*GpsB seems conformationally dynamic, possibly acting as a regulatory switch in divisome assembly.

We independently crystallized the N-terminal domain (residues 1–75) of *Sa*GpsB and determined its structure at 1.7 Å resolution (Figure 1A, Table 1). Our analysis shows an asymmetric dimer with a kinked helix conformation, in excellent agreement with the GpsB dimers seen in 8E2B.pdb (chains C/D) and 8E2C.pdb (chains A/B). Root-mean-square deviation (RMSD) between our model and 8E2B chains C/D is ~0.4 Å over all Cα atoms (70 residues), underscoring the similarity of the two structures. While 8E2B contains two *Sa*GpsB dimers in the asymmetric unit, a dimer with a kinked helix of approximately 20^°^ and a dimer with a kinked helix of approximately 40^°^, our structure best matches the 40^°^ bent helix conformation (Figure 1A). Both described *Sa*GpsB conformations vary significantly from the nearly straight helices observed in *Bs*GpsB, *Lm*GpsB, *and Sp*GpsB.

**Table 1.**
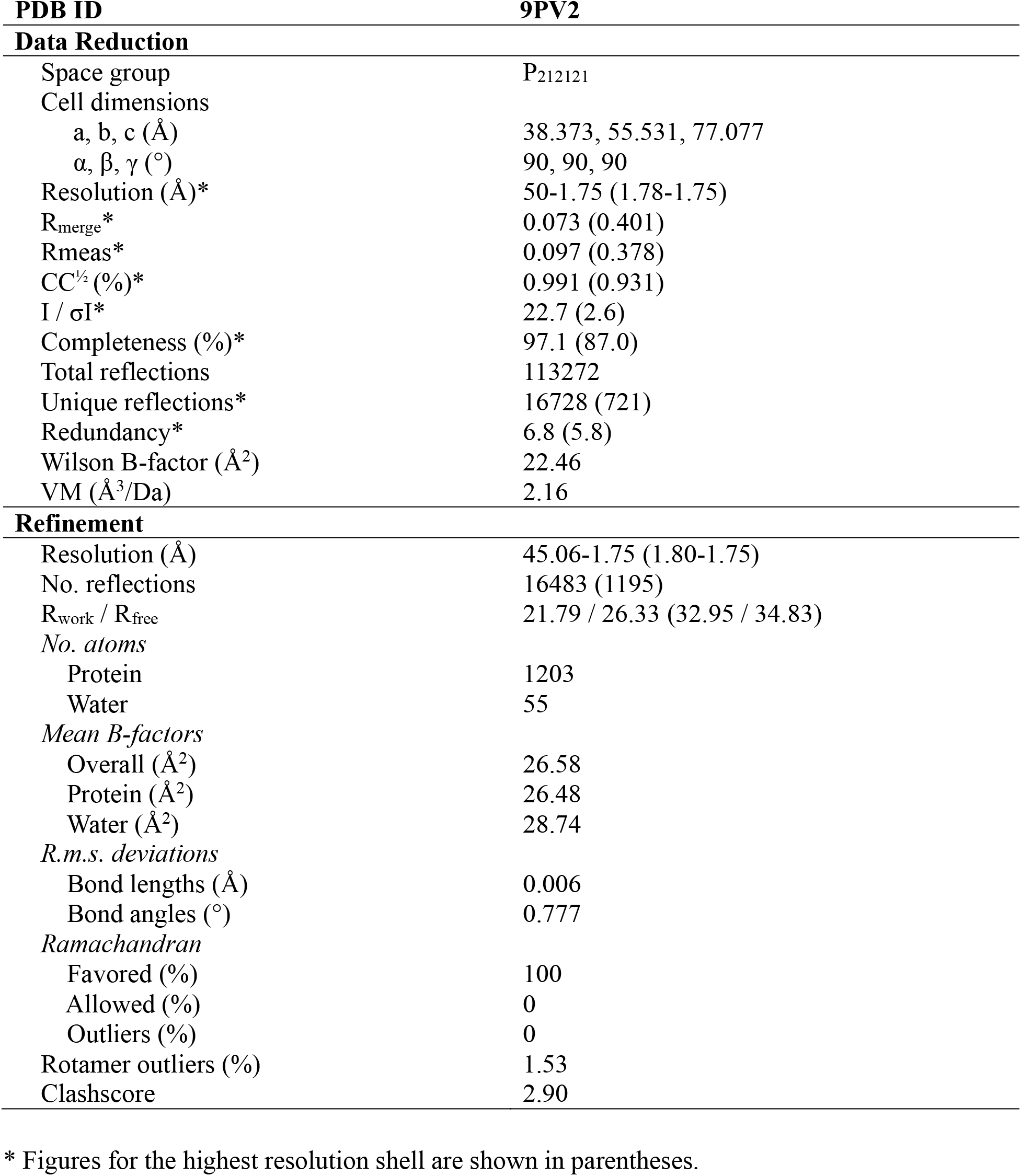
X-ray collection and refinement statistics.

**Figure 1.**
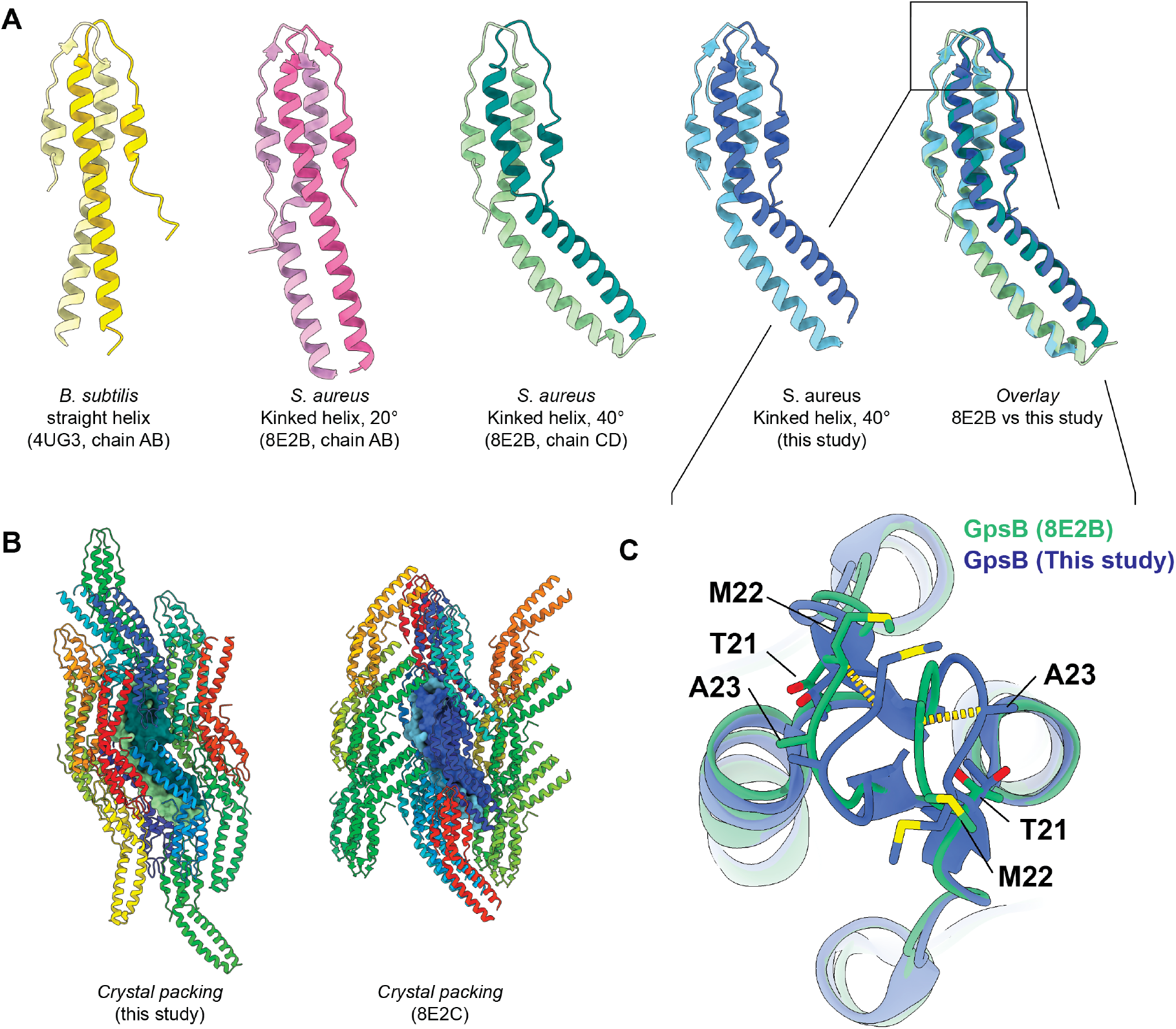
X-ray structure of *Sa*GpsB residues 1-75. **A.** Comparison of *Bs*GpsB (yellow), *Sa*GpsB conformation 1 (pink), *Sa*GpsB conformation 2 (green), and *Sa*GpsB from this study (blue). Each GpsB model is colored with two different shades of the same color to show the dimeric assembly.an overlay of the 40 kinked helix conformation from 8E2B and the 40 kinked helix structure from this study is shown (far right). **B.** Crystal packing for our *Sa*GpsB crystal structure (left) is compared with crystal packing for *Sa*GpsB + FtsZ peptide (8E3C.pdb; right). 8E2B has two copies of the GpsB dimer and was therefore not shown. While our crystal structure and 8E2C both have a single copy of GspB dimer in the asymmetric unit, their crystal packing arrangement is unique, demonstrating that the conformation observed is not influenced by the crystal lattics. **C.** Zoom-in of the membrane binding loop (aa 17-27) of the *Sa*GpsB dimer. The two conformations observed have loop displacements of 2.5-3.0 Å (yellow dashes).

Importantly, our analysis shows a unique crystal packing morphology compared to 8E2B and 8E2C (Figure 1B), confirming that the kinked-helix conformation is not an artifact of specific crystal-packing conditions. Our data therefore supports the model proposed by Sacco et al., whereby the hinge-mediated flexibility serves as a dynamic switch in *S. aureus*. Although no ligand was present in our crystals, the conformation observed is virtually identical to *Sa*GpsB bound to the (S/T)-R-X-X-R−(R/K) motif of PBP4 (8E2C.pdb), suggesting the asymmetric dimer is intrinsic to *Sa*GpsB’s fold, not induced by ligand binding. Thus, this conformation likely represents the physiologically relevant state that mediates binding to partners.

The only significant difference between our structural analysis is slight heterogeneity in the membrane-bending loop (residues ~17–27) (Figure 1C), which may reflect dynamic motion of the hinge region, as suggested in Sacco, et al. This heterogeneity is expected for a loop region proposed to insert into the inner leaflet of the membrane^1^.

Our independent crystal structure of the *Sa*GpsB N-terminal domain validates and reinforces the asymmetric, hinge-mediated conformation first described by Sacco et al. These findings support the hypothesis that hinge flexibility enables regulatory control of divisome component assembly by modulating GpsB interactions with other binding partners. Together, this work affirms GpsB’s role as a dynamic adaptor in *S. aureus* cell-division and provides a solid structural foundation for further functional studies.

## Methods

### GpsB 1-75 purification

GpsB residues 1-75 (from *S. aureus* strain NCTC 8325; Uniprot Q2FYI5; SAOUHSC_01462) were recombinantly purified as a fusion with an N-terminal 10xHis-SUMO tag. Briefly, BL21 (DE3) *E. coli* transformed with the expression plasmid were grown in Lysogeny Broth (LB) with 50 μg/mL kanamycin to mid log phase (OD_600_ ~0.6) and induced with 0.5 mM Isopropyl β-D-1-thiogalactopyranoside (IPTG) at 18 C overnight. Cells were harvested in lysis buffer (20 mM HEPES, pH 7.5; 500 mM NaCl, 20 mM Imidazole, 1 mM DTT, 1 mM PMSF), lysed by sonication, and clarified via centrifugation at 28,000 *g*. Lysate was applied to Ni+-NTA resin (GoldBio), washed with high salt buffer (20 mM HEPES, pH 7.5, 1000 mM NaCl, 20 mM Imidazole), low salt buffer (20 mM HEPES, pH 7.5, 100 mM NaCl, 20 mM Imidazole), and eluted with elution buffer (20 mM HEPES, pH 7.5, 100 mM NaCl, 300 mM Imidazole). Protein was dialyzed overnight into crystallization buffer (20 mM HEPES, pH 7.5, 100 mM NaCl, 1 mM DTT), along with SUMO protease to cleave the purification tag. The following day, the uncleaved protein and free His-SUMO tag were removed by passing the eluent over Ni+-NTA resin. The protein was further purified by anion exchange and size exclusion chromatography. The protein was concentrated to 5 mg/mL.

### GpsB 1-75 crystallization, data collection, and model building

Protein was tested for crystallization against common commercially available crystal screens using a Mosquito dropsetter (SPT Labtech) with drops composed of 200 nl protein and 200 nl reservoir solution set over 30 μl reservoir volumes. Crystals were observed within one week over a reservoir solution composed of 1.0 M Succinic Acid, 0.1 M HEPES pH 7.5, 1 %w/v PEG 2000 MME. The crystals were briefly soaked in reservoir supplemented with 15% ethylene glycol then cryocooled in liquid nitrogen. Diffraction data were collected at Southeast Regional Collaborative Access Team (SER-CAT) 22-ID beamline at the Advanced Photon Source, Argonne National Laboratory, using an incident beam of 1 Å in wavelength. Data were reduced in HKL-2000^14^. The structure was phased by molecular replacement using Phaser^15^ with PDB 8e8b as the search model^13^. Real space rebuilding were done in Coot^16^, and reciprocal space refinements and validations were done in PHENIX^17^. Coordinates and structure factors have been deposited in the Protein Data Bank (PDB) with accession number 9PV2.

## Acknowledgements

This research used resources of the Advanced Photon Source, a U.S. Department of Energy (DOE) Office of Science user facility operated for the DOE Office of Science by Argonne National Laboratory under Contract No. DE-AC02-06CH11357. SER-CAT is supported by its member institutions, equipment grants (S10_RR25528, S10_RR028976 and S10_OD027000) from the National Institutes of Health, and funding from the Georgia Research Alliance.

